# Arabidopsis NF-YCs interact with CRY2 and PIF4/5 to repress blue light-mediated hypocotyl growth

**DOI:** 10.1101/2023.04.14.536856

**Authors:** Wei Wang, Lin Gao, Tianliang Zhao, Jiamei Chen, Ting Chen, Wenxiong Lin

## Abstract

Plants can not move automatically, thus they have to perceive the changing environment (light, temperate and so on) by developing the adapted phenotypes. Light-induced hypocotyl length is an ideal phenotype for studying how plants response to light. So far, many signaling components in light-induced hypocotyl growth have been reported. Here, we focused on identifying transcription factors (TFs) involved in blue light-induced hypocotyl growth by constructing Arabidopsis TFs overexpressing lines and screening blue light-induced hypocotyl length. Finally, we found that three NF-YC proteins, NF-YC7, NF-YC5 and NF-YC8 (NF-YCs as short name), develop longer hypocotyls than the wild type under blue light. While deficient mutants, *nf-yc5nf-yc7* and *nf-yc7nf-yc8*, fail to promote the hypocotyl elongation under blue light.

NF-YCs physically interacted with CRY2 and PIF4/5, while the NF-YCs-PIF4/5 interactions were repressed by CRY2. Moreover, overexpression of CRY2 or deficiency of PIF4/5 repressed the hypocotyl elongation induced by NF-YC7 under blue light. Further investigation revealed that NF-YC7 increased the blue light-induced CRY2 degradation and regulated the activities of PIF4/5.

Taking together, this study provided a new insight into that NF-YCs function as CRY2-and PIF4/5-interacting proteins and modulate their stabilization to repress the blue light-mediated hypocotyl growth.

**Author Summary:** Light is an essential environmental factor for plants growth and development. Plants response to light signaling by displaying short hypocotyls, green leaves, and so on. The mechanisms of light responding to light have aroused extensive attention. In this study, we clarified that NF-YC family members NF-YC5/7/8 interact with CRY2, PIF4/5 and modulate their stabilization to repress the blue light-mediated hypocotyl growth. It was the first time for reporting NF-Y family members function as CRY2-and PIF4/5-interacting proteins. Therefore, this study provides a novel understanding how plants adapt to light.

## Introduction

Light, an essential and variable environmental factor, regulates the growth and development of plants. Plants have evolved adaptive phenotypes to response to variable light environment (including light quantity, quality, directionality and photoperiod). For example, seedlings in the stage of de-etiolation can produce light-induced phenotypes, such as inhibited hypocotyls, de-etiolated and expanded cotyledons when plant are exposed to light [1].

Plants sense light environment by light receptors. So far, five kinds of light receptors in Arabidopsis have been found, such as phytochromes (PHYA-E), phototropins (PHOT1 and PHOT2), zeitlupe family (ZTL, LKP, FKF), cryptochromes (CRY1 and CRY2) and UV-B receptor (UVR8) [2-4]. Among of them, CRY1 and CRY2 as blue light receptors can positively regulate blue light-induced hypocotyl growth and long day-flowering [5-7]. CRY2 mainly mediates low intensity blue light-induced photomorphogenesis [5]. CRY2 can be activated by light-dependent dimerization and phosphorylation [8-10]. Finally, light-activated CRY2 is degraded via ubiquitination [11-13].

In the CRYs-meidated photomorphogenesis, many signaling partners, such as bHLH family members PIFs (Phytochrome Interacting Factors), can interact with CRYs and regulate the expressions and activities of light-responsive genes [14-16]. PIF4 and PIF5 as the light-response repressors are necessary for limiting blue light-induced hypocotyl growth by directly interacting with CRY1 and CRY2 [14]. Moreover, CRY2 and PIF4/5 associate with the same promoter regions to regulate PIFs targets [14].

*PIF4/5* deficient mutants *pif4* and *pif5* showed no obvious hypocotyl growth compared with the wild type under limiting blue light [14], while the sustained activation of PIF lead to the hypocotyl over elongation. Therefore, the regulation of PIFs activities is necessary for promoting photomorphogenesis. PIFs may be inactivated by the transcriptional regulation or the post-translational modification [17]. For the transcriptional regulation, phytochromes can inhibit the activities of PIFs by preventing them from binding to their targets [18, 19]. For the post-translational modification, PIFs are inactivated by protein kinases through rapid phosphorylation prior to consequent ubiquitin-mediated degradation [18-23].

As previously reported, NF-YB1-NF-YC12 heterodimer form a complex with bHLH144, a member of bHLH family, to regulate the grain quality in rice [24]. Except for that, several Arabidopsis NF-YC members were identified as the positive regulators of photomorphogenesis, such as NF-YC1, NF-YC3, NF-YC4 and NF-YC9. These four NF-YC proteins redundantly promote photomorphogenesis by light-dependently interacting with HDA15 (Histone Deacetylase 15), modulating histone H4 acetylation level and co-repressing the expressions of hypocotyl elongation-related targets [25]. In addition, NF-YC1, NF-YC3, NF-YC4 and NF-YC9 inhibit BR synthesis and interact with BIN2 to suppress BR signal pathway during light-induced hypocotyl growth [26].

In *Arabidopsis*, NF-YC family consists of 13 members [27, 28]. As mentioned above, previous reports mainly claimed how NF-Y factors regulate light-induced hypocotyl growth pathway as positive regulators [25, 26, 29]. In this study, we demonstrated that three Arabidopsis NF-YC proteins, NF-YC5, NF-YC7 and NF-YC8 (NF-YCs as the short name), inhibit blue light-inhibited hypocotyl elongation. NF-YCs directly interact with CRY2, PIF4 and PIF5 to promote CRY2 degradation and regulate PIF4/5 activities. CRY2 inhibits NF-YCs-PIFs interactions. This work will be helpful for understanding how NF-YCs suppress blue light-mediated hypocotyl growth.

## Materials and Methods

### Plant Materials

All the mutants used in this study were in the background of Arabidopsis Columbia ecotype. Mutants *nf-yc5-1* (salk_130605), *nf-yc7-1* (salk_012179), *nf-yc8-1* (cs862600) and *nf-yc8-2* (salk_064020) were obtained from ABRC (the Arabidopsis Biological Resource Center, https://www.arabidopsis.org). Mutants *pif4*, *pif5* and *pif4pif5* were kindly provided by Dr. Ying Li (Yangzhou University, China). Mutants *nf-yc5-2* (inserted “A” at 455 bp) and *nf-yc7-2* (inserted “T” at 229 bp) were produced using Crispr (Clustered regularly interspaced short palindromic repeats, **S1 Fig**) and confirmed by sequencing. To prepare double mutant *nf-yc5nf-yc7*, *NF-YC5* was mutated under the background of *nf-yc7-1* using Crispr because of the close physical locations of *NF-YC5* and *NF-YC7* in the same chromosome. Double mutant *nf-yc7nf-yc8* was generated by crossing *nf-yc7-1* and *nf-yc8-1*.

To generate Arabidopsis TFs overexpressing lines FGFP-TFs, more than 1000 transcription factors in Arabidopsis pENTR/D-TF Collection (Stock CD4-88, ABRC) were cloned into the expression vector *pACT2::FGFPbar* (FGFP: short name of <u>F</u>lag and <u>GFP</u>-infused protein) using LR recombination reaction (Invitrogen). To prepare CRY2 overexpressing line Myc-CRY2, the coding sequences of *CRY2* amplified from Arabidopsis Columbia ecotypes was infused into *pACT2::cMYChyg* using the In-fusion method (Transgen, China). All above plasmids were transformed into *rdr6-11* (RNA dependent polymerase 6 mutant that can repress gene silence) by *Agrobacterium tumefaciens*–mediated floral-dip method [30-32]. Plant expression vectors *pACT2::cMychyg* and *pACT2::FGFPbar* used in this study were modified from *pCambia3301*. All the primers used in this study were listed in S1 Table.

The co-overexpression line Myc-CRY2/FGFP-NF-YC7 was obtained by crossing Myc-CRY2 with FGFP-NF-YC7. To check the genetic relationships between NF-YC7 and PIF4/5, the coding sequence of *NF-YC7* was infused into *pACT2::cMychyg* and introduced into *Col-0* and *pif4pif5,* respectively.

### Growth Conditions

Seeds were planted on MS medium, kept at 4 ℃ under dark for 4 days, and then treated under white light for 1 day at 21 ℃. After treated by the above processes, seeds were grown in intelligent growth chamber at 21 ℃ with different light treatment according to the experiment plans. The blue light source and the green light source were from Haibo Instrument and Equipment Company (Changzhou, China). The illumination meter used in this study is TES-1339P (TES, China).

### Hypocotyl Length Measurement

To measure the hypocotyl length, all the seeds were treated using continuous blue light (10 μmol m^−2^ sec^−1^) or dark for 6 days. At least 20 seedlings of each sample were measured using Image J. Data was analyzed using Excel and GraphPad Prime. The experiments were repeated three times at least and the results of one repeat were shown in this study.

### GUS Staining

To check the expression patterns of *NF-YCs*, *NF-YCs* promoters (*NF-YCspro*) as described were amplified by high-fidelity PCR[27]. *NF-YCs* promoters were respectively infused into the 5’ terminal part of FGUS (<u>F</u>lag and <u>GUS</u> fused protein) in the expression vector *pFGUSbar* derived from the binary vector *pCambia3301*. *NF-YCspro::FGUS* were introduced into *Col-0*. After treated under dark, blue light or long day (16 h light/ 8 h dark) for 6 days, the transgenic seedlings carrying *NF-YCspro::FGUS* were collected and incubated in GUS staining buffer in the dark for overnight at 37 ℃, and then washed using absolute ethanol till the chlorophyll totally disappeared as described [33]. Pictures were taken using Nikon SMZ18.

### Degradation of CRY2

To verify the roles of NF-YCs in the CRY2 degradation, we employed two groups of genotypic materials. FGFP-NF-YC7 and the relative wild type (the transgenic line carrying *pACT2::FGFPbar*) were used to check the effect of NF-YC7 on the endogenous CRY2 degradation. In addition, Myc-CRY2 and Myc-CRY2/FGFP-NF-YC7 were used to analyze the effect of NF-YC7 on the CRY2 degradation. All the above indicated seedlings grown under dark for 6 days were exposed to blue light (10 μmol m^−2^ sec^−1^) for 0, 5, 10, 20 minutes with or without 50 μM MG132 (Selleck, USA) incubation. Protein extraction and detection were performed as previously reported [11, 34]. CRY2 were detected using Anti-CRY2 antibody (Abcam, UK).

### Co-IP Assay

For co-IP assay in *Arabidopsis*, Myc-CRY2/FGFP (the line co-overexpressing CRY2 and *pACT2::FGFPbar* empty vector) and Myc-CRY2/FGFP-NF-YCs were employed to check the effects of light on the NF-YCs-CRY2 interactions in *Arabidopsis.* All the seedlings grown under dark for 8 days were exposed to blue light (0, 5, 10, 15 min) and collected respectively. Then, samples treated under blue light for 5, 10 and 15 minutes were mixed together to make sure that the time of light treatment was enough and CRY2 was still not degraded fully. The previous described protocol of protein extraction and purification with minor modifications was followed in this study [11].

Co-IP assay in HEK293T cells was performed to identify *in vitro* protein interactions. To verify the NF-YCs-CRY2 interactions, GFP, GFP-NF-YCs and Myc-CRY2 driven by CMV5 promoter were expressed in HEK293T cells according to the plans. To analyze the NF-YCs-CRY2-PIFs interactions, GFP-NF-YCs, Myc-CRY2 and Flag-PIFs driven by CMV5 promoter were expressed in HEK293T cells according to the plans.

Cells carrying the indicated plasmids were treated under dark or blue light (100 μmol m^−2^ sec^−1^, 1 h) and collected. The previous described protocols of HEK293T cells transformation and protein extractions were followed [11].

Protein complex was purified using GFP-Trap agarose beads (Chromotek) and separated by western blot. Anti-GFP antibody (MBL), anti-Myc antibody (MBL) and anti-Flag antibody (MBL) were used to detect FGFP/GFP-NF-YCs, Myc-CRY2 and Flag-PIFs, respectively.

### Firefly Luciferase Complementation Imaging Assay (LCI)

For LCI assay, the coding sequences of *NF-YCs* were fused into the N-terminal of nLUC and the coding cequences of *PIFs* were cloned into the C-terminal of cLUC as described [35]. To check the interactions between NF-YCs and PIFs, NF-YCs-nLUC or cLUC-PIFs were transiently expressed into *N. benthamiana*. To check the effect of CRY2 on NF-YCs-PIFs interactions, NF-YCs-nLUC and cLUC-PIFs were transiently expressed into *N. benthamiana*, in the absence or presence of Myc-CRY2. The protocol of transient expression were followed as previous described [36]. After transformated for two days, the *N. benthamiana* leaves expressing the nLUC or cLUC-fused proteins were incubated with 1 mM luciferin under dark for 5 minutes. LUC singals were detected using In Vivo Plant Imaging System (Night Shade LB 985).

### Dynamic Expressions of PIFs

To generate the PIFs-overexpressing lines Flag-PIF4/5 and the co-overexpression lines GFP-NF-YC7/Flag-PIF4/5, the coding sequences *NF-YC7*, *PIF4* and *PIF5* were infused into *pACT2::GFPsul* or *pACT2::Flagbar* modified from *pCambia3301* and introduced into *rdr6-11*. To check the expression dynamics of PIF4/5 regulated by NF-YC7, the transgenic seedlings grown under dark for 6 days were exposed to blue light (10 μmol m^−2^ sec^−1^) for 0, 1, 24, 48, 72 h. Samples were collected into liquid nitrogen immediately. Protein was extracted and separated by western blot. Flag-PIF4 and Flag-PIF5 were detected using anti-Flag antibody (MBL). Anti-GFP antibody was used to detect GFP-NF-YC7. Non-specific bands were used as the loading control.

### Accession Numbers

RDR6 (AT3G49500), NF-YC5 (AT5G50490), NF-YC7 (AT5G50470), NF-YC8 (AT5G27910), CRY2 (AT1G04400), PIF4 (AT2G43010), PIF5 (AT3G59060).

## Results

### NF-YCs redundantly and negatively regulate blue light-mediated hypocotyl growth

To identify TFs that regulate blue light-induced photomorphogenesis, we respectively overexpressed about 1000 Arabidopsis TFs in *rdr6-11,* and then screened hypocotyl lengths of TFs-overexpressing lines FGFP-TFs under dark or blue light (10 μmol m^−2^ sec^−1^). Finally, we found that two independent NF-YC7-overexpressing lines, FGFP-NF-YC7-175 and FGFP-NF-YC7-176, showed longer hypocotyls than the wild type *rdr6-11* under blue light, but no differences under dark (**Fig 1A, D**). To investigate the role of two close NF-YC7 homologs NF-YC5 and NF-YC8 in blue light-induced photomorphogenesis, we analyzed hypocotyl lengths of their overexpression lines under dark and blue light (**Fig 1B, C, E, F**). Similarly, the hypocotyl lengths of NF-YC5-or NF-YC8-overexpressing lines, FGFP-NF-YC5-s3, FGFP-NF-YC5-85, FGFP-NF-YC8-79 and FGFP-NF-YC8-80, were longer than that of the wild type *rdr6-11* under blue light, while no difference under dark, which means NF-YC5, NF-YC7 and NF-YC8 may redundantly repress blue light-induced hypocotyl growth. To further identify this hypothesis, we employed *NF-YCs* single mutants (*nf-yc5-1*, *nf-yc5-2*, *nf-yc7-1*, *nf-yc7-2*, *nf-yc8-1 and nf-yc8-2*) and double mutants (*nf-yc5nf-yc7* and *nf-yc7nf-yc8*), and examined the blue light-induced hypocotyl phenotypes (**Fig 1G-J**). The hypocotyl lengths of all the single mutants showed no difference from that of the wild type *Col-0* under both blue light and dark conditions (**Fig 1G, I**). As expected, double mutants *nf-yc5nf-yc7* and *nf-yc7nf-yc8* showed shorter hypocotyls under blue light, but similar to the wild type *Col-0* under dark (**Fig 1H, J**). Consequently, NF-YC7, NF-YC5 and NF-YC8 redundantly and negatively regulate the blue light-mediated hypocotyl growth.

**Fig 1.**
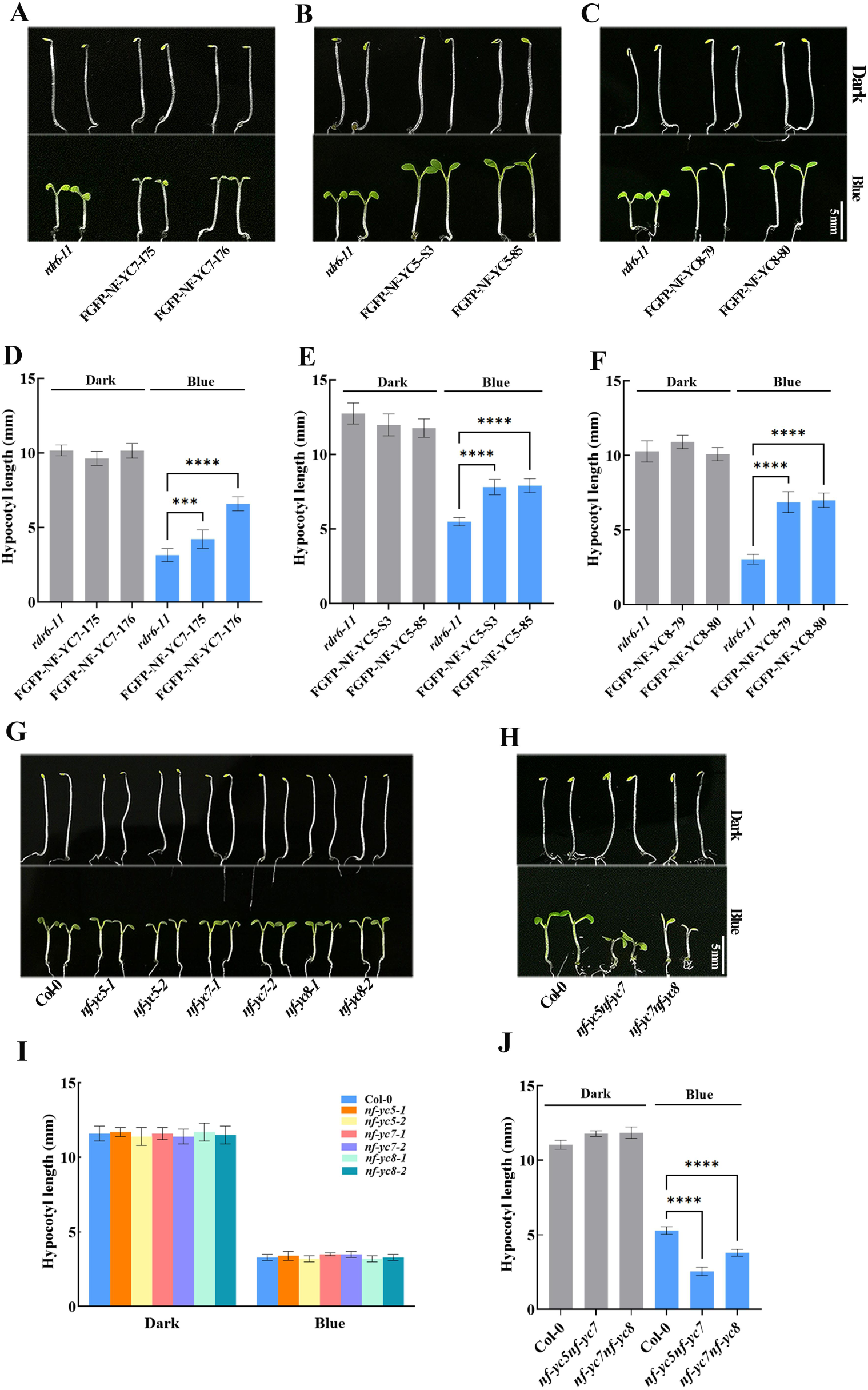
NF-YCs negatively regulate blue light-meidated hypocotyl growth. (**A-C**) The representative hypocotyl images of the wild type (*rdr6-11*), two independent NF-YCs-overexpressing lines (FGFP-NF-YC7-175, FGFP-NF-YC7-176, FGFP-NF-YC5-s3, FGFP-NF-YC5-85, FGFP-NF-YC8-79 and FGFP-NF-YC8-80). (**D-F**) Hypocotyl length of the seedlings shown in (**A-C**). (**G-H**) The representative hypocotyl images of the wild type (Col-0), single mutants (*nf-yc5-1*, *nf-yc5-2*, *nf-yc7-1*, *nf-yc7-2*, *nf-yc8-1* and *nf-yc8-2*) and double mutants (*nf-yc5nf-yc7* and *nf-yc7nf-yc8*). (**I-J**) Hypocotyl length of the seedlings shown in (**G-H**). All the indicated seedlings grown under blue light (10 μmol m^-2^ s^-1^) or dark for 6 days were used to measure hypocotyl length (***P<0.001, ****P<0.0001, compared with the wild type grown under blue light; one-way ANOVA; ±SD, n=20). All experiments were repeated at least three times and the data from one repeat was shown.

### Light induces the dynamical expressions of NF-YCs in hypocotyl

As the repressors of the blue light-mediated hypocotyl growth, the expressions of NF-YCs in hypocotyl may be regulated by light. To check this assumption, we performed GUS staining assay (**Fig 2A**). The results showed that the main localization of FGUS was in the apical or basal regions of hypocotyls when the transgenic seedlings carrying *NF-YCspro::FGUS* were grown under dark for 6 days. FGUS was accumulated in hypocotyls when the seedlings were grown under blue light for 6 days. Especially, FGUS was detected in the whole hypocotyls when the seedlings were exposed to white light (16 h light/ 8 h dark) for 6 days. Moreover, we tested the FGUS expression under dark, blue light or white light by western blot (**Fig 2B**). Accordance with the above results, FGUS was increased under blue light and white light, compared with that under dark. All the results revealed NF-YCs promoters are light-responsive promoters and light induces the accumulation of NF-YCs in hypocotyl.

**Fig 2.**
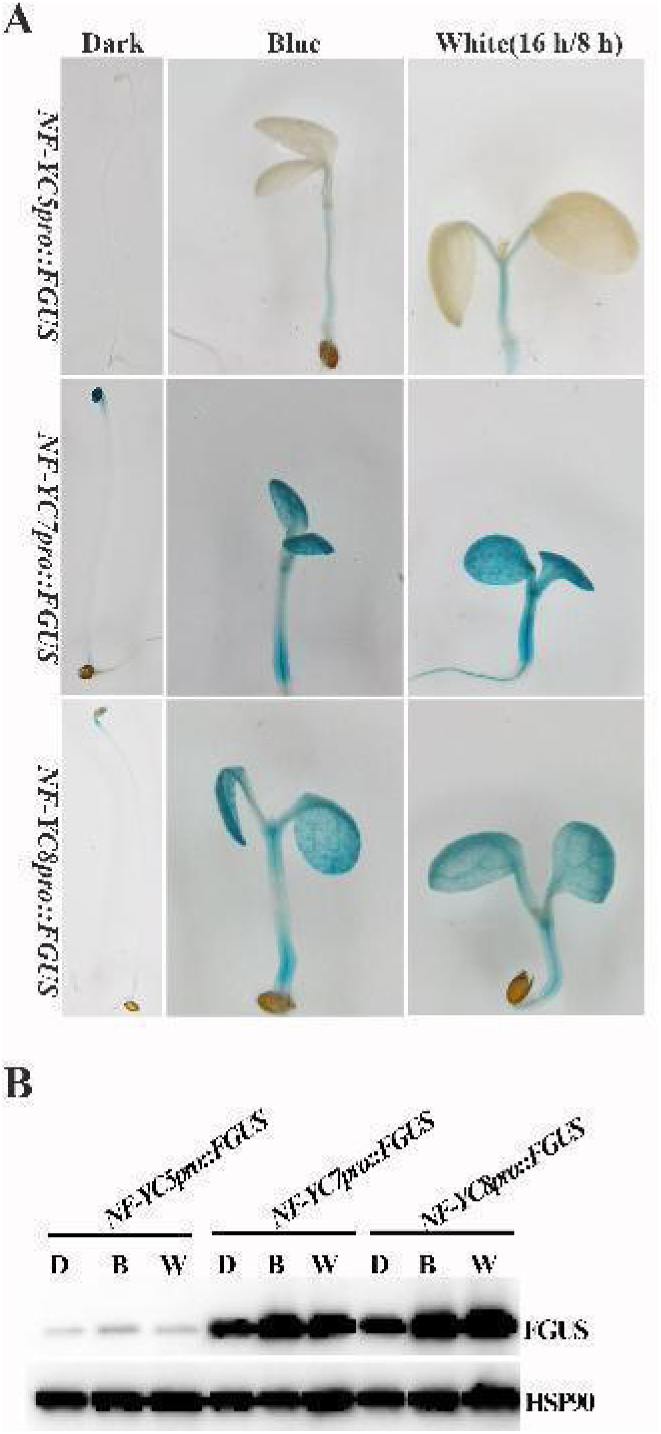
Light induces NF-YCs expressions in hypocotyl. (**A**) GUS staining of the transgenic seedlings carrying *NF-YCspro::FGUS* grown under dark, continuous blue light (10 μmol m^-2^ s^-1^) or white light (16 h light/ 8 h dark) for 6 days. (**B**) The expression levels of FGUS in the seedlings shown in (**A**). FGUS was probed by anti-Flag antibody. The loading control was probed by anti-HSP90.

### NF-YCs directly interact with CRY2

It was reported that many molecules (such as CRY2, BIC1/2, HY5, and so on) regulate blue light-induced hypocotyl growth [5, 9, 37]. Whether do NF-YCs interact with them to regulate blue light-mediated hypocotyl growth? We employed co-IP assays in HEK293T cells and Arabidopsis and conformed that NF-YC7 directly interacts with CRY2 (**S3 Fig**). Furthermore, we identified the interactions between NF-YCs and CRY2. Myc-CRY2 was pulled by NF-YCs in HEK293T cells under both dark and blue light (**Fig 3A**), and NF-YCs-CRY2 interactions were further confirmed in *Arabidopsis* (**Fig 3B**). Among tested NF-YCs, NF-YC7 showed the strongest interaction with CRY2 in *Arabidopsis*. Together, NF-YCs directly interact with CRY2 in a blue light-independent manner.

**Fig 3.**
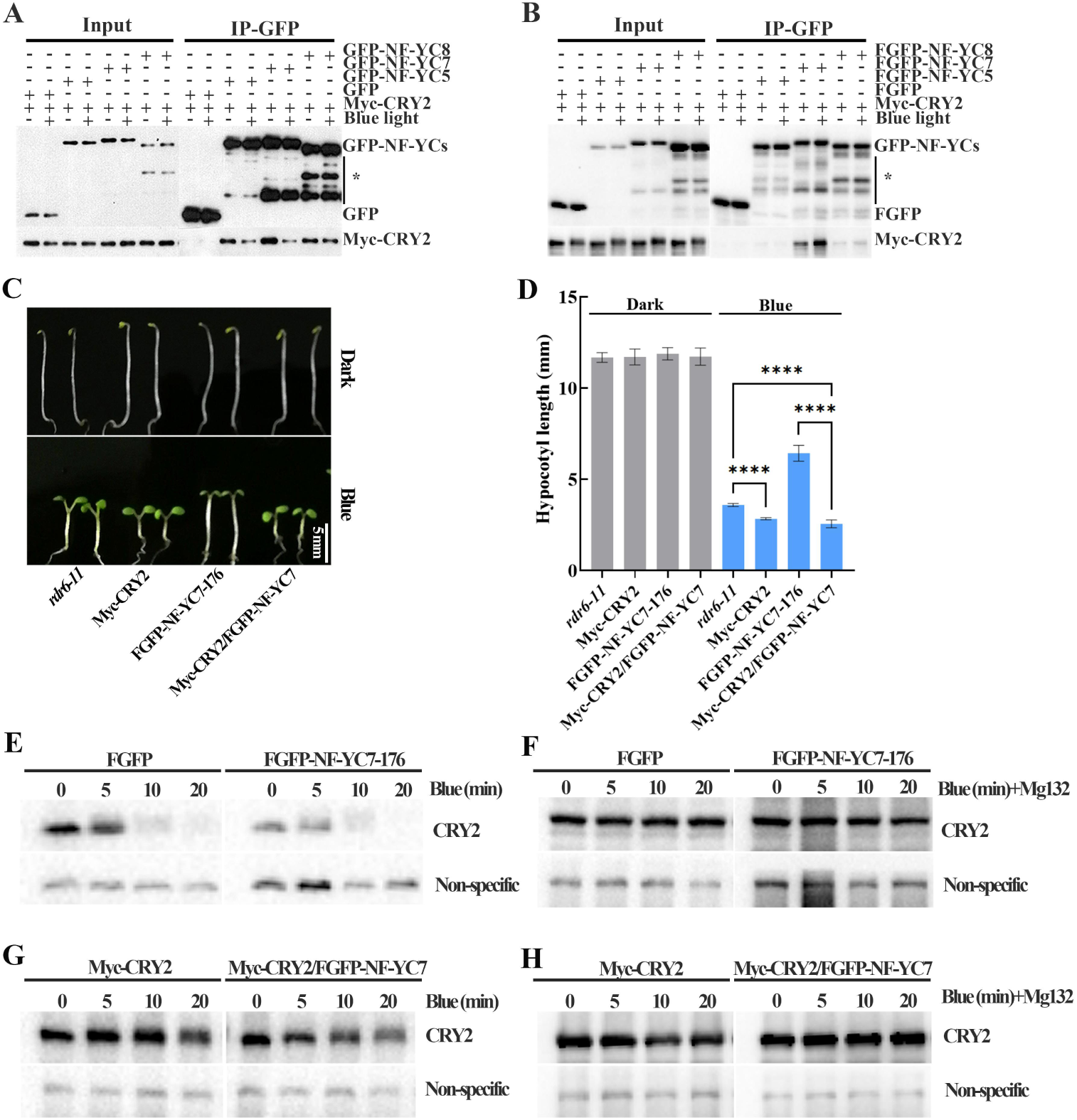
NF-YC7 negatively regulates CRY2-mediated hypocotyl growth under blue light. **(A-B)** The NF-YCs-CRY2 interactions *In vitro* **(A**) *and In vivo* **(B**). The HEK293T cells expressing Myc-CRY2/GFP or Myc-CRY2/GFP-NF-YCs treated by blue light (100 μmol m^-2^ s^-1^) for 1 h or dark were used to check the CRY2-NF-YCs interactions (**A**). The co-overexpression lines Myc-CRY2/FGFP and Myc-CRY2/FGFP-NF-YCs grown under blue light (10 μmol m^-2^ s^-1^) or dark for 6 days were used to check the CRY2-NF-YCs interactions in *Arabidopsis* (**B**). Protein extracts were immunoprecipitated by GFP agarose beads. The IP signal or the co-IP signal was probed by anti-GFP or anti-Myc antibodies, respectively. Star means non-specific bands. (**C-D**) The function of NF-YC7 in CRY2-meiated hypocotyl growth. The representative images of the wild type (*rdr6-11*), CRY2-overexpressing line Myc-CRY2, NF-YC7-overexpressing line FGFP-NF-YC7-176 and the co-overexpression line Myc-CRY2/FGFP-NF-YC7 (**C**). (**D**) Hypocotyl length of the indicated genotypes shown in (**C**). The above-mentioned seedlings were grown under blue light (10 μmol m^-2^ s^-1^) or dark for 6 days (****P<0.0001; one-way ANOVA; ± SD, n=20). All experiments were repeated at least three times and the data from one repeat was shown. (**E-F**) NF-YC7 regulates the CRY2 degradation. (**E**) The endogenous CRY2 degradation in the relative wild type (FGFP-overexpressing line in *rdr6-11*) and FGFP-NF-YC7-176. (**G**) The degradation of CRY2 in Myc-CRY2 and Myc-CRY2/FGFP-NF-YC7. All the indicated seedlings grown in the dark for 6 days were exposed to blue light (10 μmol m^-2^ s^-1^) for 0, 5, 10 and 20 min with or without Mg132 incubation. CRY2 was detected using anti-CRY2 antibody. Non-specific bands were used as the loading control.

### NF-YC7 negatively regulate the CRY2-mediated hypocotyl growth by inactivating CRY2

Based on the physical interactions between NF-YCs and CRY2 (**Fig 3A-B**), we detected the genetic relationships between NF-YC7 and CRY2 in blue light-mediated hypocotyl growth. Hypocotyl lengths of Myc-CRY2, FGFP-NF-YC7, Myc-CRY2/FGPF-NF-YC7 and the wild type were examined under dark and blue light (**Fig 3C-D**). There were no difference among the tested seedlings under dark. Blue light induced shorter hypocotyl in Myc-CRY2 and a longer hypocotyl in FGFP-NF-YC7, compared with the wild type. Myc-CRY2/FGFP-NF-YC7 showed similar hypocotyl length to Myc-CRY2, shorter than the wild type under blue light. CRY2 repressed blue light-induced hypocotyl elongation induced by FGFP-NF-YC7. That means CRY2 is the downstream of NF-YC7 in blue light-inhibited hypocotyl elongation.

The hypocotyl length of NF-YC7-overexpressing lines was similar to that of *cry2-1* under blue light, longer than that of the wild type. Therefore, we purposed that NF-YC7 might inactivate CRY2, and tested whether NF-YC7 effects the degradation of CRY2 in *Arabidopsis* (**Fig 3E-H**). The endogenous CRY2 began to degrade not only in FGFP-NF-YC7 but also in control plants (FGFP-overexpressing line), since the etiolated seedlings grown under dark for 6 days were exposed to blue light. The endogenous CRY2 degradation increased with the extension of blue light treatment. However, endogenous CRY2 degraded in FGFP-NF-YC7 faster than the wild type (**Fig 3E**). Consistently, CRY2 degradation was faster in Myc-CRY2/FGFP-NF-YC7 than Myc-CRY2 under blue light (**Fig 3F**). Because of the requirement of blue light-dependent polyubiquitination for CRY2 degradation [12], we analyzed the effect of NF-YC7 on CRY2 degradation pretreated with MG132 (proteasome inhibitor). After the incubation of MG132, the blue light-induced CRY2 degradation was fully defected in the present of or the absent of NF-YC7 (**Fig 3F and H**). Moreover, our initial results verified that NF-YCs interact with E3 ubiquitin ligases, such as COP1, LRB1 and LRB2 using LCI assay (**S4 Fig**). That suggested that NF-YC7 may increase the degradation of CRY2 by polyubiquitination.

### NF-YCs interact with CRY2, PIF4 and PIF5

As reported, CRY2 interacted with PIF4/5 directly (Pedmale *et al*., 2016). We hypothesized that NF-YCs might interact with PIF4/5 as well. Using the LCI assay (**Fig 4A**), NF-YCs-nLUC and cLUC-PIF4/5 were found to interact with each other (**Fig 4A**). Further, we checked the effects of CRY2 on the NF-YCs-PIF4/5 interactions. NF-YCs-nLUC and cLUC-PIF4/5 were co-expressed with or without Myc-CRY2 in tobacco leaves (*N. benthamiana*). The LUC signal of NF-YCs-nLUC and cLUC-PIF4/5 was decreased by Myc-CRY2 (**Fig 4A**). In HEK293T cells, PIF4/5 were pulled by NF-YCs, which was dramatically disturbed by Myc-CRY2 (**Fig 4B**). NF-YCs directly interact with PIF4/5, and the interactions was suppressed by CRY2.

**Fig 4.**
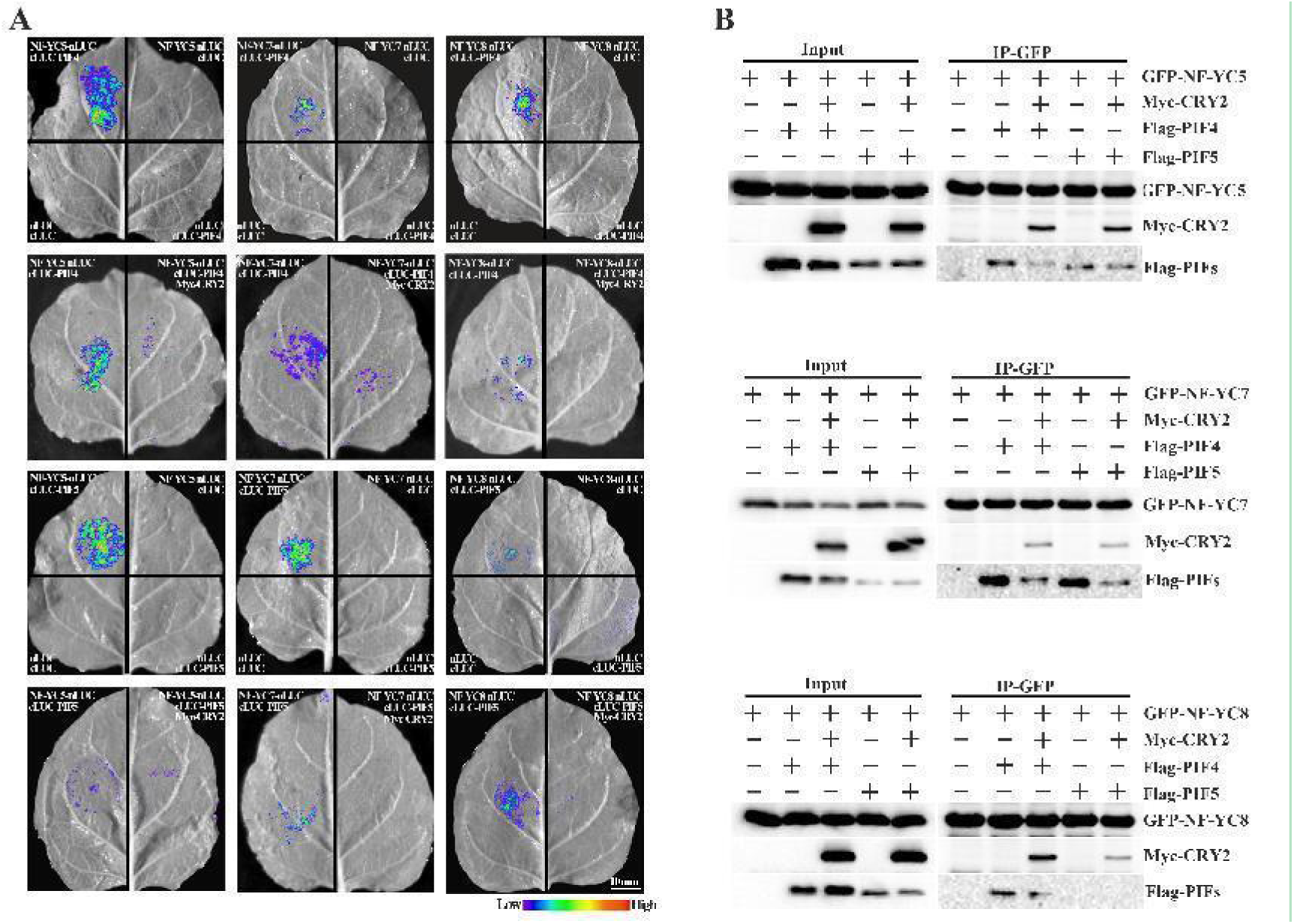
The interactions between NF-YCs, CRY2 and PIFs. (**A**) The interactions between NF-YCs, CRY2 and PIFs using LCI assay in *Tobacco*. For NF-YCs-PIFs interactions, *Agrobacterium* strains carrying the indicated plasmids pairs were transiently expressed in Tobacco leaves as shown, respectively. One *Tobacco* leaf was evenly divided into 4 parts: 1, NF-YCs-nLUC and cLUC-PIFs were co-expressed in the left upper part; 2, NF-YCs-nLUC and cLUC were co-expressed in the right upper part; 3, nLUC and cLUC-PIFs were co-expressed in the right lower part; 4, nLUC and cLUC were co-expressed in the left lower part. Except for the 1^st^ group, the other 3 groups were the negative controls. To check that CRY2 effects NF-YCs-PIFs interactions in *Tobacco*. One *Tobacco* leaf was divided into two parts along the main leaf vein. NF-YCs-nLUC and cLUC-PIFs were transiently co-expressed in the left part of *Tobacco* leaf. Myc-CRY2, NF-YCs-nLUC and cLUC-PIFs were transiently co-expressed in the right part of the same *Tobacco* leaf. LUC signal was visualized after the incubation of Luciferin under dark for 5 min. The exposure time was equal to 5 min. (**B**) The interactions between NF-YCs, CRY2 and PIFs using co-IP assay in HEK293T cells. GFP-NF-YCs were co-expressed with Flag-PIFs or Myc-CRY2 in HEK293T cells according to the indicated plans. GFP-NF-YCs were immunoprecipitated using GFP agarose beads. The IP signal (GFP-NF-YCs) and the co-IP signal (Myc-CRY2, Flag-PIFs) were detected by anti-GFP, anti-Myc and anti-Flag antibodies, respectively. Star means non-specific bands. Plus and minus mean that the candidate proteins are present and absent.

### PIF4 and PIF5 function as the downstreams of NF-YCs

PIF4 and PIF5 are responsible for blue light-mediated hypocotyl elongation [38]. Consistent with above report, we showed that deficient mutants of *PIF4* and *PIF5* result in shorter hypocotyl, compared with the wild type grown under blue light for 6 days (**S5 Fig**). To investigate whether PIF4/5 participate in NF-YCs-regulated hypocotyl growth, the hypocotyl phenotypes of the seedlings *pif4pif5,* Myc-NF-YC7, and Myc-NF-YC7/*pif4pif5* were observed under dark and blue light (**Fig 5C-D**). After grown under dark for 6 days, all the above genotypes lead to similar hypocotyl lengths. Compared with that of the wild type, longer hypocotyl of Myc-NF-YC7 and shorter hypocotyl of *pif4pif5* were exhibited after the treatment of blue light. The hypocotyl length of Myc-NF-YC7/*pif4pif5* were similar to that of *pif4pif5,* shorter than that of the wild type under blue light. Conclusively, PIF4/5 are the downstreams of NF-YC7 in blue light-induced hypocotyl growth.

**Fig 5.**
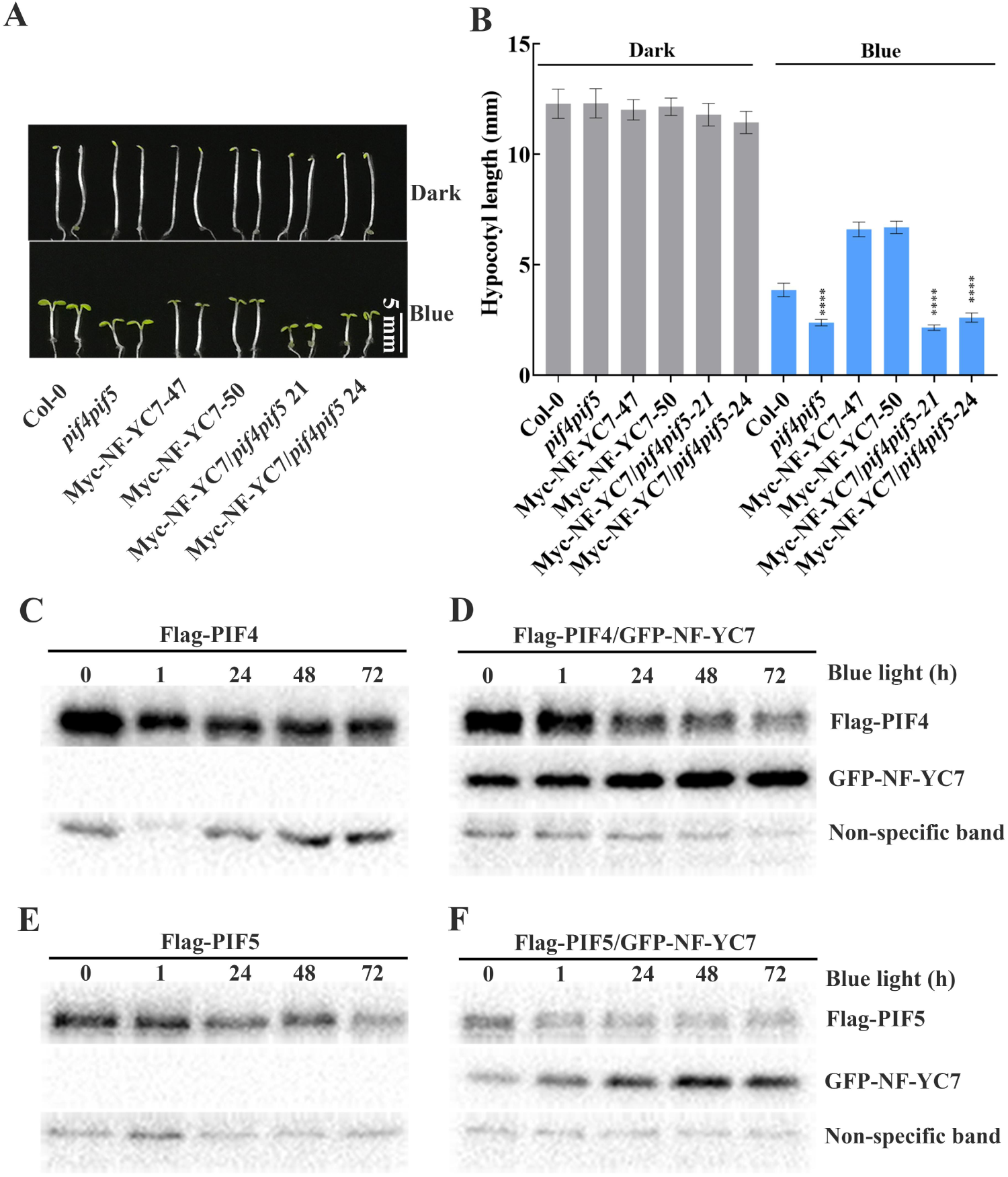
PIFs are the downstream of NF-YC7 in the blue light-inhibited hypocotyl elongation. (**A-B**) The representative hypocotyl images (**A**) and hypocotyl length (**B**) of Col-0, *pif4pif5*, two independent NF-YC7-overexpressing lines (Myc-NF-YC7-47 and Myc-NF-YC7-50) and two independent lines overexpressing NF-YC7 in *pif4pif5* (Myc-NF-YC7/*pif4pif5*-21, Myc-NF-YC7/*pif4pif5*-24). All the seedlings for measuring hypocotyl length were grown under blue light (10 μmol m^-2^ s^-1^) or dark for days (****P<0.0001, compared with the wild type grown under blue light; one-way ANOVA; ±SD,n=20). All experiments were repeated at least three times and the data from one repeat was shown. (**E-F**) The effect of NF-YC7 on the activities of PIF4 (**E**) and PIF5 (**F**). Flag-PIF4/5: PIFs-overexpressing line in *rdr6-11*, Flag-PIF4/GFP-NF-YC7 and Flag-PIF5/GFP-NF-YC7: the co-overexpression lines of PIF4/5 and NF-YC7 in *rdr6-11*. All the seedlings grown under dark for 6 days were exposed to blue light (10 μmol m^-2^ s^-1^) for 0, 1, 12, 24, 72 h and collected to check the PIF4/5 activities using western blot. GFP-NF-YC7 and Flag-PIF4/5 were detected by anti-GFP and anti-Flag antibodies, respectively. Non-specific bands were the loading controls.

To check how NF-YCs regulate PIF4/5, we examined the effect of NF-YC7 on PIF4/5 on the protein level, using Flag-PIFs (PIF4/5-overexpressing lines) and Flag-PIFs/GFP-NF-YC7 (the line co-overexpressing of PIF4/5 and NF-YC7). PIF4 and PIF5 were slightly degraded when all the seedlings grown under dark for 6 days were exposed to blue light (**Fig5E-F**). However, the migration speed of PIF4/5 in Flag-PIFs/GFP-NF-YC7 was slower than that in Flag-PIFs when seedlings grown under dark were exposed to blue light. That means NF-YC7 may regulate PIF4/5 activities in blue light-inhibited hypocotyl elongation.

## Discussion

Arabidopsis NF-YC family contains 13 homologs (NF-YC1 to NF-YC13). It has been reported that NF-YC1, NF-YC3, NF-YC4 and NF-YC9 play redundant and positive roles in light inhibition of hypocotyl elongation by light-dependently interacting with HDA15 and modulating histone H4 acetylation level [25, 26, 29]. And these four NF-YC proteins integrate the BR and light signaling pathways by interacting with BIN2 to control light-induced hypocotyl growth [26]. In this study, we demonstrated another three Arabidopsis NF-YC proteins, NF-YC7, NF-YC5 and NF-YC8, redundantly and negatively regulate blue light inhibition of hypocotyl elongation by physically interacting with CRY2 and PIF4/5 and regulating their activities (**Fig 6**). That means NF-YC family encodes two above clades of NF-YC proteins, which play an opposite role in photomorphogenesis, to contribute to moderate hypocotyl growth under light.

**Fig 6.**
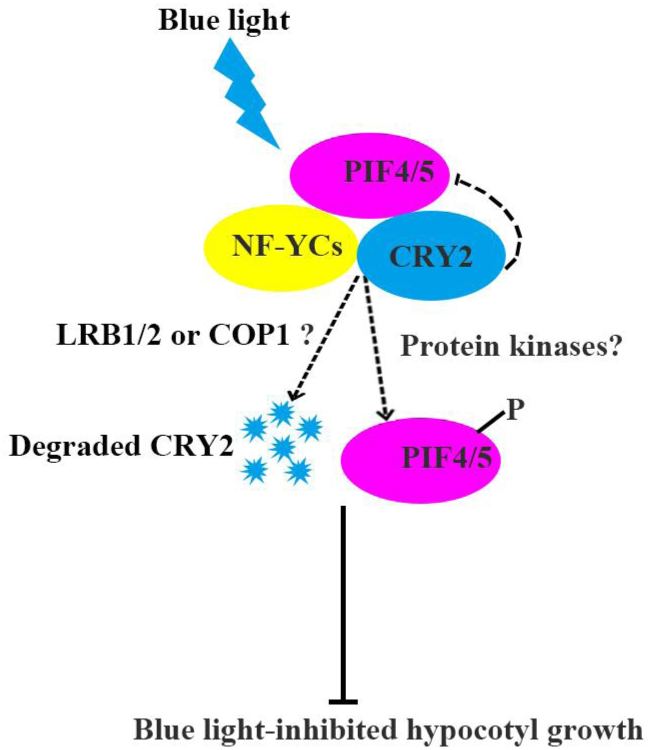
A hypothetic model depicting blue light-mediated hypocotyl growth. Based on this model, NF-YCs interact with CRY2 and PIF4/5 upon blue light exposure. However, the interactions between NF-YCs and PIF4/5 were weaken in the present of CRY2. NF-YCs increase CRY2 degradation and keep PIF4/5 phosphorylation to repress blue light-mediated hypocotyl growth.

As the suppressors of blue light-inhibited hypocotyl growth, NF-YCs are localized in hypocotyl and increased in light-dependent manner (**Fig 5E-F**). However, Siefers and colleagues reported that the localizations of NF-YC1, NF-YC3, NF-YC4, NF-YC9, NF-YC10, NF-YC11, and NF-YC12 are in hypocotyl using GUS staining assay, not including NF-YC5, NF-YC7 and NF-YC8 [27]. Obviously, we showed the different expression patterns of NF-YC5, NF-YC7 and NF-YC8 in hypocotyl from the above report [27]. We reasoned that the endogenous NF-YCs are low and their expressions are induced by light. That is why we checked the expression patterns of NF-YCs in dark, blue light and white light conditions.

NF-YCs physically interact with CRY2 in blue light-inhibition of hypocotyl growth (**Fig 3A-B**). Actually, it was the first proof that NF-YCs act as CRY2-interacting proteins. A handful of CRY2 interacting proteins have been reported to regulate the photoactivation and inactivation of CRY2. PPKs (Photoregulatory Protein Kinases 1, 2, 3 and 4) phosphorylate dimeric CRY2 to enhance CRY2 activity [8]. For CRY2 inactivation, photo-excited CRY2 is polyubiquitinated by LBR1/2 (Light-Response Bric-a-Brack/Tramtrack/Broad 1 and 2) and degraded by the 26s proteasome [11]. BIC1 (Blue-light Inhibitor of Cryptochromes 1) interacts with CRY2 to suppress CRY2 dimerization and the interactions between CRY2 and its signaling partners [9]. In our study, NF-YCs act as CRY2-interacting partners to promote CRY2 degradation. Additionally, our preliminary result showed that NF-YCs interact with CRY2-related E3 ligases, COP1 and LRB1/2 (**S4 Fig**). Hence, NF-YCs-E3 ligases interactions (**S4 Fig**) and the light-induced accumulation of NF-YCs (**Fig 2A-B**) provide a compete explanation for NF-YCs recruiting the indicated E3 ligases or enhance their activities to accelerate CRY2 degradation.

CRY2 directly interacts with PIF4/5 and modulates PIFs activities in limiting blue light-mediated photomorphogenesis [14]. Our results showed that NF-YCs can directly interact with PIF4/5 and CRY2 (Fig4A-B). However, the interactions between NF-YCs and PIF4/5 weakened or disappeared in the present of CRY2. That suggested that CRY2 might compete NF-YCs with PIFs or deactivate PIFs.

PIFs stabilization is light intensity-dependent [39]. Limiting blue light can keep PIF4 protein constant, but promote the accumulation of PIF5, while high intensity light can rapidly decline the abundance of bHLH members, such as PIF1, PIF3, PIF4 and PIF5 [14, 40]. Blue light intensity used in our study lead to the slight PIF4/5 degradation (**Fig 5E-F**). As phosphorylation proteins, specific phosphorylation status of PIF4/5 triggered by light can be followed by proteasome-mediated degradation [41]. Our results showed that the migration of PIF4/5 was slower in the present of NF-YC7 (**Fig 5E-F**). That means that NF-YC7 may regulate PIF4/5 activities by keeping their phosphorylation status. Some known protein kinases, such as BIN2 (BRASSINOSTEROID-INSENSITIVE 2), can interact with NF-YC1/3/4/9 and phosphorylate PIF4 [26, 42]. It is possible that NF-YC7 interact with some known or unknown protein kinases to phosphorylate and destabilize PIFs.

Summarily, we proposed that NF-YCs suppress blue light-inhibited hypocotyl elongation by interacting with CRY2 and PIF4/5 to increase CRY2 degradation and modulate PIF4/5 activities (**Fig 6**). The mechanism of regulation and interaction of NF-YCs and hypocotyl growth-related signaling partners are complicated.

## Acknowledgements

We thank Dr. Ying Li (Yangzhou University, China) for kindly providing PIF4/5 deficient mutants *pif4, pif5* and *pif4pif5*, Dr. Yuling Chen (Hebei Normal University, China), Dr. Wenfei Wang (Agriculture and Forestry University, China) and Dr. Qiang Zhu (Fujian Agriculture and Forestry University, China) for helpful suggestions, and Dr. Libo Han (Fujian Agriculture and Forestry University, China) and Dr. Yadi Chen (Yangzhou University, China) for technical support with LCI assay.

## Funds

This work was supported by the National Science Foundation of China (http://www.nsfc.gov.cn) (Grant 31701241) to WW, the Natural Science Foundation of Fujian Province (http://www.fjkjt.gov.cn) (Grant 2018J05045) to WW and the China Postdoctoral Science Foundation (http://www.chinapostdoctor.org.cn) (Grant 2020M682062) to WW. The funders had no role in study design, data collection and analysis, decision to publish, or preparation of the manuscript.

## Author contributions

W.W conceived research plans, carried out the experiments, analyzed the data and wrote the manuscript; W.X.L supervised the experiments and modified the manuscript; L.G, T.L.Z, J.M.C and T.C participated in this project. All authors have approved the manuscript and declared that no competing interests exist.

## Supplemental Materials

S1 Fig Information and identification of *NF-YC*s-related genetic materials S2 Fig FGFP-NF-YC7 driven by ACT2 promoter is located in hypocotyl S3 Fig NF-YC7 interact with CRY2 in HEK293 cells

S4 Fig NF-YCs interact with CRY2-related E3 ligases

S5 Fig PIF4/5 modulate blue light-induced hypocotyl growth S1 Table Primers in this study

S2 Table Primary and Secondary antibodies used in this study

## Notes

### Competing Interest Statement

The authors have declared that no competing interests exist.

